# Towards a more accurate quasi-static approximation of the electric potential for neurostimulation with kilohertz-frequency sources^*^

**DOI:** 10.1101/2023.08.25.554885

**Authors:** Thomas Caussade, Esteban Paduro, Matías Courdurier, Eduardo Cerpa, Warren M. Grill, Leonel E. Medina

**Affiliations:** Instituto de Ingeniería Matemática y Computacional, Facultad de Matemáticas, Pontificia Universidad Católica de Chile, Santiago, Chile; Departamento de Matemática, Facultad de Matemáticas, Pontificia Universidad Católica de Chile, Santiago, Chile; Department of Biomedical Engineering, Department of Electrical and Computer Engineering, Department of Neurobiology, Department of Neurosurgery, Duke University, Durham, NC, USA; Departamento de Ingeniería Informática, Universidad de Santiago de Chile, Santiago, Chile

**Keywords:** neurostimulation, quasistatic approximation, Helmholtz, kilohertz-frequency signals

## Abstract

**Objective:** Our goal was to determine the conditions for which a more precise calculation of the electric potential than the quasi-static approximation may be needed in models of electrical neurostimulation, particularly for signals with kilohertz-frequency components.

**Approach:** We conducted a comprehensive quantitative study of the differences in nerve fiber activation and conduction block when using the quasi-static and Helmholtz approximations for the electric potential in a model of electrical neurostimulation.

**Main results:** We first show that the potentials generated by sources of unbalanced pulses exhibit different transients as compared to those of energy-balanced pulses, and this is disregarded by the quasi-static assumption. Secondly, the relative errors for current-distance curves were below 3%, while for strength-duration curves the variations ranged between 1-17%, but could be improved to less than 3% across the range of pulse duration by providing a corrected quasi-static conductivity. Third, we extended our analysis to trains of pulses and reported a “congruence area” below 700 Hz, where the fidelity of fiber responses is maximal for suprathreshold stimulation. Further examination of waveforms and polarities revealed similar fidelities in the congruence area, but significant differences were observed beyond this area. However, the spike-train distance revealed differences in activation patterns when comparing the response generated by each model. Finally, in simulations of conduction-block, we found that block thresholds exhibited errors above 20% for repetition rates above 10 kHz. Yet, employing a corrected value of the conductivity improved the agreement between models, with errors no greater than 8%.

**Significance:** Our results emphasize that the quasi-static approximation cannot be naively extended to electrical stimulation with high-frequency components, and notable differences can be observed in activation patterns. As well, we introduce a methodology to obtain more precise model responses using the quasi-static approach, which can be a valuable resource in computational neuroengineering.

## 1 Introduction

Electrical stimulation of nerve fibers or other neural elements is used to treat the symptoms of myriad neural disorders and diseases such as Parkinson’s disease, chronic pain, epilepsy, and others [17]. Electrical currents are applied to the tissues by means of implanted or surface electrodes and a pulse generator that delivers pulses at a repetition rate of typically no greater than 200 Hz, and up to a few kHz in cochlear implants [31] and spinal cord stimulation [15]. Electrical stimulation is commonly modeled using a two-step approach in which, by assuming that the presence of the neurons does not affect the electric field, the electric field generated by an external electrode is calculated, and then, the resulting potentials are applied to a cable model of a nerve fiber [11, 19].

For electric field calculation, most modeling studies rely on the quasi-static assumptions and simplify Helmholtz’s equation to Laplace’s equation [28]. Under the quasi-static assumption, capacitive, inductive, and propagation effects are neglected, and in general, all three conditions are met if the frequency of the stimulation signal is sufficiently low. However, if the frequency content of the electric field is too high, then the quasi-static assumption may not be appropriate [5, 22, 14, 33]. For example, in a homogeneous, isotropic volume conductor with a point-source electrode, the error between thresholds calculated using the quasi-static approximation and the solution of Helmholtz’s equation averaged about 13% for current pulses of 0.025 – 1 ms [5]. In addition, the error in potentials calculated with the quasi-static assumption is time-dependent, peaking at the edges of the pulse, and therefore during repetitive stimulation, the error may vary depending on the pulse repetition rate.

Further, kilohertz-frequency (KHF) signals have received increased attention due to the potential to block the conduction of nerve impulses [16, 4, 21]. This interest has manifested in the use of KHF signals, for example, in applications of epidural spinal cord stimulation for chronic pain relief, and transcutaneous tibial stimulation for pain management [32, 23]. Analyses of these applications may require to use the appropriate equations to obtain accurate calculations of the electric field due to KHF sources.

We quantified the responses of model nerve fibers to point-source stimulation, after calculation of the electric field using either the quasi-static assumption or Helmholtz’s equation. We hypothesized that fiber responses can significantly differ between these solutions, especially for KHF stimulation. We estimated the error of the quasi-static assumption under various conditions, including single-pulse stimulation, and pulse-train stimulation for a wide range of repetition rates. Our study builds upon previous work that determined the conditions under which the quasi-static assumption may not be appropriate for single pulse stimulation [5]. We extended this analysis to include nerve fiber activation, and conduction block during stimulation with pulse trains. Further, we introduce a novel methodology to determine a value of the conductivity that leads to smaller errors in model fiber responses when using the quasi-static approximation, thereby improving model estimation. The results have implications for neurostimulation modeling and design of electroceutical devices, particularly in applications that use KHF signals [1, 35]

## 2 Background

We used a two-step approach to quantify nerve fiber responses to electrical stimulation. First, we calculated the electric potentials generated by a point-source electrode in an infinite, homogeneous, isotropic, and dispersive volume, and then we applied these potentials to a cable model of a nerve fiber. The electric properties of the medium can be represented by the complex permittivity *ε*_*c*_, defined as

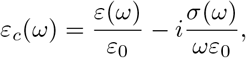

where *ε* and *σ* are the permittivity and conductivity of the medium, respectively, and depend on the frequency, *ω*, of an externally applied source. Furthermore, living tissues do not exhibit relevant magnetic response [2, 12], so we set the magnetic permeability *μ* as the free-space permeability *μ*_0_ = 4*π*·10^−7^*H/m*.

We note that the transmembrane potential, defined as the difference between extracellular and intracellular potentials, may be affected by the presence of the fiber itself [18], but here we neglect this effect.

### 2.1 Electric potentials

We seek a closed-form expression of the time-harmonic electric potential, *ϕ*. For practical purposes, all the subsequent potentials should be regarded as the voltage between any point of interest and infinity, where we set the reference potential to zero. The time-harmonic potential can be calculated by solving the inhomogeneous Helmholtz (IH) equation [28], given a current density **J** := **J**(***r***) generated, in this case, by a point source placed at ***r***_0_ ∈ ℝ^3^ away from the fiber. The scalar Helmholtz’s equation (using the time convention *e*_^*iωt*^_) is

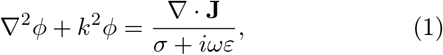

where 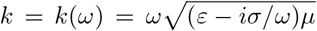 is the complex propagation constant. Invoking Green’s representation theorem, the solution evaluated at any point in space distinct from ***r***_0_ can be expressed as

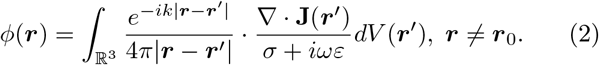

Given the spherical symmetry of the setup, an analytical solution of (2) can be obtained in terms of the source current intensity, *I*, by simply evaluating the integral:

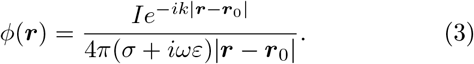

The quasi-static (QS) approximation assumes that the capacitive, inductive, and propagation effects can be neglected. The first assumption states *σ* ≫ *ωε*, thereby neglecting the contribution of the imaginary part to the potential in the denominator (3) associated with the capacitive response. The second assumption neglects any magnetic contribution to the electric field. Finally, the third assumption simplifies the complex exponential term of the propagation effects by assuming that |*k*| *·* |***r***−***r***_0_| ≪ 1. The resulting potential corresponds to the so-called *quasi-static potential* :

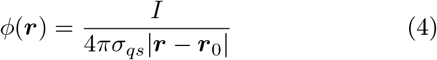

where the conductivity *σ*_*qs*_ represents a “quasi-static conductivity” that does not depend on the stimulation frequency and is the effective conductivity assigned to the medium. A thorough comparison and validity of both models for single monophasic pulses can be found in [5]. In this work, we extended this analysis to compare the responses to trains of biphasic pulses, not only for fiber activation (spike trains), but also for conduction block.

### 2.2 Dispersive dielectric properties

Living tissues typically exhibit dispersive behavior, *i*.*e*., their responses to electric fields depend on the frequency of the field [13]. The frequency-dependent permittivity and conductivity are computed from the complex permittivity, *ε*_*c*_, using multiple Cole-Cole dispersions [8] at low, medium, high, and very-high frequencies. The resulting model defines the complex permittivity as [13]:

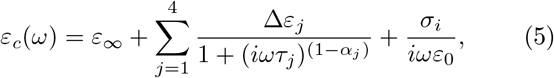

where *ε*_∞_ is the permittivity as *ω* → ∞, *σ*_*i*_ is the ionic conductivity, Δ*ε*_*j*_ corresponds to the difference between static and high-frequency limit permittivities, *τ*_*j*_ is the time-relaxation constant, *α*_*j*_ is the distribution parameter as a result of adjusting the spectral shape. Hence, for a given angular frequency, *ω*, the frequency-dependent parameters are obtained as

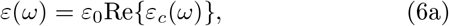

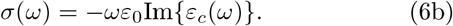

In this work, we used the parameters for brain grey matter [13], and calculated the conductivity and permittivity using (6), from the complex permittivity defined in (5).

### 2.3 Time-dependent potentials

Consider now the time-dependent current density **J**_*s*_ generated by the point source located at ***r***_0_, and a periodic shape-function *s*(*t*) with an associated fundamental frequency *ω*:

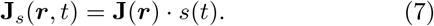

To include transient effects, we set the fundamental frequency according to the total simulation time *T*, so that *T* = 2*π/ω*. All stimulus effects are expected to occur in a finite time window, so we set the interval [0, *T*] large enough to contain the period of interest.

The shape function, *s*(*t*), can be represented using a Fourier series. Let {*a*_*m*_}_*m*∈ℤ_ be the complex Fourier coefficients, which can be directly computed for a specific stimu-lus waveform:

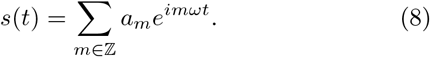

To include the dispersive dielectric properties described in Section 2.2, we solve the IH equation (1) for each harmonic *mω* individually, with the right-hand side scaled by the corresponding coefficient *a*_*m*_. Assuming that the medium is linear, we applied superposition over each term. Next, to obtain the time-dependent full-spectrum response *ϕ*_*ih*_, each harmonic’s response is multiplied by its corresponding Fourier basis term, yielding

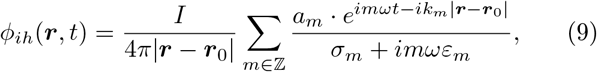

where *k*_*m*_ := *k*(*mω*), *ε*_*m*_ := *ε*(*mω*), *σ*_*m*_ := *σ*(*mω*) are computed accordingly for the *m*-th harmonic. To derive the QS solution *ϕ*_*qs*_, we recall the same assumptions used to obtain (4):

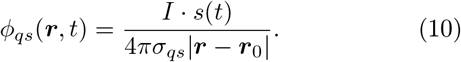

### 2.4 Quasi-static conductivity

We note that in the calculation of *ϕ*_*qs*_, selecting *σ*_*qs*_ may be arbitrary, and it is unclear what value to draw from the dispersion curve, *e*.*g*., conductivity at dc, at the repetition rate or something in between. Previous work suggested that the QS approximation is valid in applications of transcranial current stimulation for frequencies of up to ∼2 MHz [14], and that the error of the QS in electromagnetic field calculation in a model of the human head is less than 5% for frequencies up to 25 MHz [24]. Yet, total neglect of the permittivity further narrows this range of validity [14]. There is evidence that the brain’s tissue conductivity can be estimated by solving Maxwell equations [7], and more recent work proposes to use a representative frequency-specific value of the conductivity at 1 kHz, also adapted to local electric field intensity, for electroconvulsive therapy [33]. Here, we propose a more general framework to determine a *σ*_*qs*_ value that provides QS potentials closer to those of the whole IH equation. Conduction block can be predicted by average delivered power [27], and for the range of pulse durations considered in this work, the required power for fiber activation appears to be barely affected by the stimulation waveform [36]. Thus we present a power-based argument that we tested for biphasic rectangular pulses, but may be used for any waveform shape.

In our approach, we matched the average apparent power delivered to the tissue, particularly at the center of the fiber, by each model to compute the effective conductivity that should be employed in the QS model. Consider the root mean square (RMS) voltage as

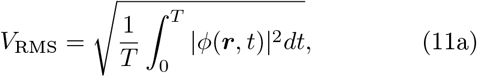

for any ***r*** ∈ ℝ^3^, and RMS current

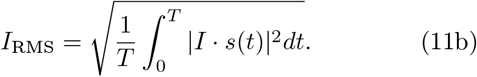

Then, the average apparent power transferred to ***r*** is given by the product *V*_RMS_ · *I*_RMS_. Since the *I*_RMS_ does not depend on the chosen electrical field model, and imposing that the average power delivered to the fiber be the same for QS and IH, it follows that:

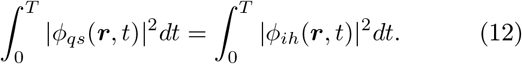

Therefore, from equations (10) and (9), and using Parseval’s identity, we define the *corrected quasi-static conductivity* 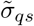 as:

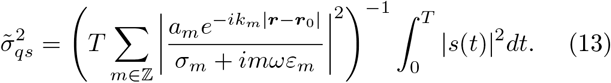

We remark there is an explicit dependence on the distance to the source, given by the propagation effects, requiring this new value to be space-dependent. For simplicity, we fixed |***r*** − ***r***_0_| as the distance from the source to the fiber, as the fiber section closest to the point-source electrode is more likely to be affected by the external potential, at least for cathodic stimulation [29].

## 3 Methods

### 3.1 Fiber model

We used the MRG double cable model of a myelinated nerve fiber [19], implemented in Neuron 8.2 [6]. We used model fibers of diameters of 7.3, 10, 12.8, and 16 *μ*m, with 84 nodes of Ranvier, resulting in long enough fibers to avoid end effects.

For all simulations, we set the temperature at 37 ^◦^C and the initial membrane voltage at -77.3 mV. The model was solved using implicit Euler integration with time steps of 0.1 – 1 *μ*s.

### 3.2 Stimulation waveform

Consider the stimulation with *N* pulses repeated with frequency *f*_*r*_ (*i*.*e*., period *T*_*r*_ = 1*/f*_*r*_) starting at *t* = *p*_*s*_. Then let *p*_*n*_ := *p*_*s*_ + *n · T*_*r*_ for *n* = 0, 1,… *N* − 1. We modeled various trains of pulses, with pulse width Δ and cathodic polarity (the negative phase comes first). Consider the symmetric *biphasic* pulse train

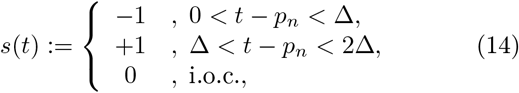

We refer to *full-duty cycle* trains as symmetric biphasic pulse trains that verify the limit case 2Δ · *f*_*r*_ = 1, *e*.*g*., trains of pulses where half-cycle is anodic and half-cycle is cathodic while active.

For all simulations, the external electric potential generated by a point-source electrode above the middle of the fiber was computed using (10) for the QS assumption, or (9) for the exact solution of the IH equation. For the latter, we obtained the Fourier coefficients of the waveform using equation (14), in a closed-form expression that we evaluated directly for each harmonic, for a frequency content of at least 500 kHz. For conduction block experiments, we employed up to 2 MHz content to ensure roughly 50 harmonics for each pulse. When needed, to determine the corrected effective conductivity we computed numerically 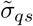 in (13), using the corresponding waveform as defined above and their respective Fourier coefficients. However, due to the periodic nature of signals, we deduced the RMS voltages in (12) do not depend on the number of pulses, so this parameter is irrelevant to the determination of 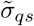,Then we simplified the calculation by computing the Fourier coefficients of the waveform with *N* = 1. In other words, repetitive or single-pulse stimulation may be performed with the same corrected value.

### 3.3 Activation thresholds

We used single-pulse extracellular stimulation with a point-source electrode to compute strength-duration (SD) and current-distance (IX) curves, for different fiber diameters. For SD simulations, the pulse duration was varied from 5 *μ*s to 500 *μ*s, with a fixed electrode-to-fiber distance of 1 mm. For the IX curves, we varied the electrode-to-fiber distance from 0.05 mm to 5 mm, while keeping the pulse duration constant at 100 *μ*s. For each of these setups, we computed activation thresholds (±0.01 nA), *i*.*e*., minimum amplitude to generate an action potential using a single pulse, and verified that an action potential propagated across the fiber at 40 nodes of Ranvier from the middle node.

Let *I*_*qs*_ and *I*_*ih*_ be the activation threshold current required by the QS and IH potentials, respectively. We quantified the percent error between the thresholds estimates as a function of pulse duration and distance for SD and IX, respectively, as:

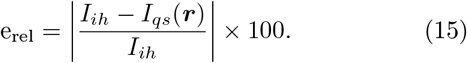

The stimulus was initiated after *p*_*s*_ = 10 ms, and we stopped the simulation after 5 ms to ensure the return to baseline.

### 3.4 Repetitive pulse stimulation

We analyzed the activation patterns and conduction block responses of MRG fibers to extracellular stimulation using trains of biphasic pulses, for the potentials calculated with the QS assumption and the exact IH solution. The source is placed at 1 mm distance from the center of the fiber.

To analyze the activation of spike trains, we considered trains of pulses with a fixed pulse duration of 100 *μ*s and repetition rates between 100 Hz and 2 kHz, starting at 10 ms, then active for a period of 40 ms, and finally ending after an additional 10 ms. We computed the fidelity index, *i*.*e*., the number of action potentials divided by the number of applied pulses and multiplied by a factor of 100, for amplitudes varying from 0.0 mA to 0.9 mA. Furthermore, let fid*_QS_* and fid*_IH_* be the reported fidelity using the QS approximation and the exact IH solution respectively. We define the fidelity error as

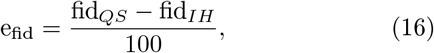

noting the absence of absolute value, in order to interpret which model had the highest fidelity, given a particular set of parameters.

In addition, we calculated the spike-train distance [34], which essentially returns the minimal cost to transform a spike train into another. In brief, this is performed by adding or deleting spikes (cost 1 for every modified spike) and translations of Δ*t* in time (with cost *q*Δ*t*). In the same original work, an efficient algorithm to determine the minimal-cost path of operations is proposed. Here, *q* is an arbitrary non-negative parameter. We note that for *q* = 0 the spike-train distance accounts for the difference in the number of spikes or action potentials, while for large values of *q*, it can be interpreted as a measure of dissimilarity between trains of pulses.

Finally, for the conduction block experiments, we considered extracellularly applied trains of pulses with repetition rates ranging from 4 to 40 kHz. We computed block thresholds as the minimal current amplitude required to block the propagation of action potentials along the fiber. So, we applied a supra-threshold single intracellular pulse of amplitude 10 nA and duration 30 *μ*s at one end of the fiber, 40 ms after the extracellular stimulus started (see [3, figure1]). Then, to test for conduction block, we measured the voltage at the other end of the fiber and verified the propagation or not of the test action potential, and, block thresholds were obtained after a binary search (±0.1 *μ*A). We repeated the experiments for trains with fixed pulse duration at 10 *μ*s and with full-duty cycle trains.

**Figure 1:**
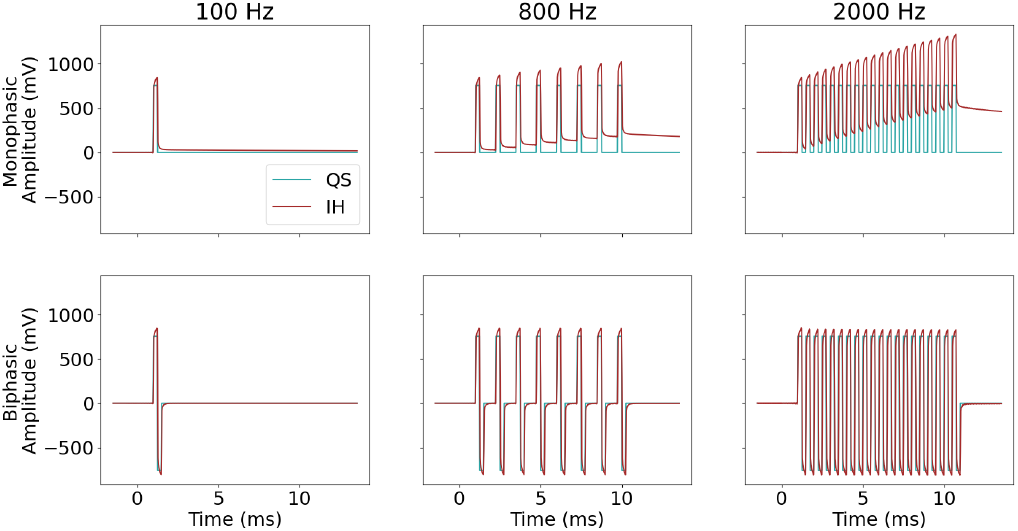
Trains of monophasic (top row) and biphasic (bottom row) rectangular pulses of 100 *μ*s with different repetition rates (columns). The source is active for 10 ms and begins at 1 ms. The electric potential is computed using (10) for the quasi-static model (QS) with *σ*_*qs*_ = 0.105 S/m, and (9) for the inhomogeneous Helmholtz’s solution (IH).

## 4 Results

We compared model fiber responses to extracellular potentials calculated with the QS approximation with fixed conductivity *σ*_*qs*_ = 0.105 S/m, with corrected quasi-static conductivity given by the definition (13), and with the exact solution to the IH equation.

### 4.1 Transient effects

We explored the transient effects due to the signal’s energy balance. Consider the illustrative example in figure 1, where we computed electric potentials using equations (9) and (10) for trains of monophasic and biphasic pulses. To have sufficient simulation time to visualize transient effects and to enforce the Fourier coefficient *a*_0_ to be small, independently from the shape, we set *T* = 200 ms, resulting in a fundamental frequency of 5 Hz.

As suggested by equation (10), the QS solution is simply a scaled version of the source’s waveform by the quasi-static conductivity, current amplitude, and distance. In contrast, the IH exact solution accounts for capacitive effects that prevent abrupt variations (continuous yet exponentially fast), and propagation effects. As a consequence, the timing between signals to reach their maximum and minimum amplitudes is slightly delayed.

The potentials differed, and this difference was accentuated for unbalanced trains of pulses. For monophasic pulses, as the train is narrowed by increasing the repetition rate, the interpulse intervals are insufficient for the potential to return to baseline, thus resulting in a gradual accumulation of charge over time. However, there are no detectable time-dependent differences between the QS and IH potentials for trains of balanced biphasic pulses.

### 4.2 Excitation properties

We calculated activation thresholds for MRG fibers with various fiber diameters, using single biphasic pulses for both the QS and IH models. We replicated experiments in [5, figure 6] for single monophasic pulses.

The errors between activation thresholds did not exceed 2% for distances of 0.05–5 mm (figure 2a), and ranged from less than 1% to 17% for a pulse duration of 5–500 *μ*s (figure 2b). These results indicate that there is good agreement between the QS and IH models, provided there is an appropriate choice of conductivity, consistent with the previous results in [5]. Consequently, we recomputed the SD curves using the same setup but employing the corrected quasistatic conductivity (figure 2c). Errors were *<* 3% for all pulse durations, thus demonstrating that the QS model can be adjusted to match better the fiber responses obtained with the exact solution of the IH equation.

**Figure 2:**
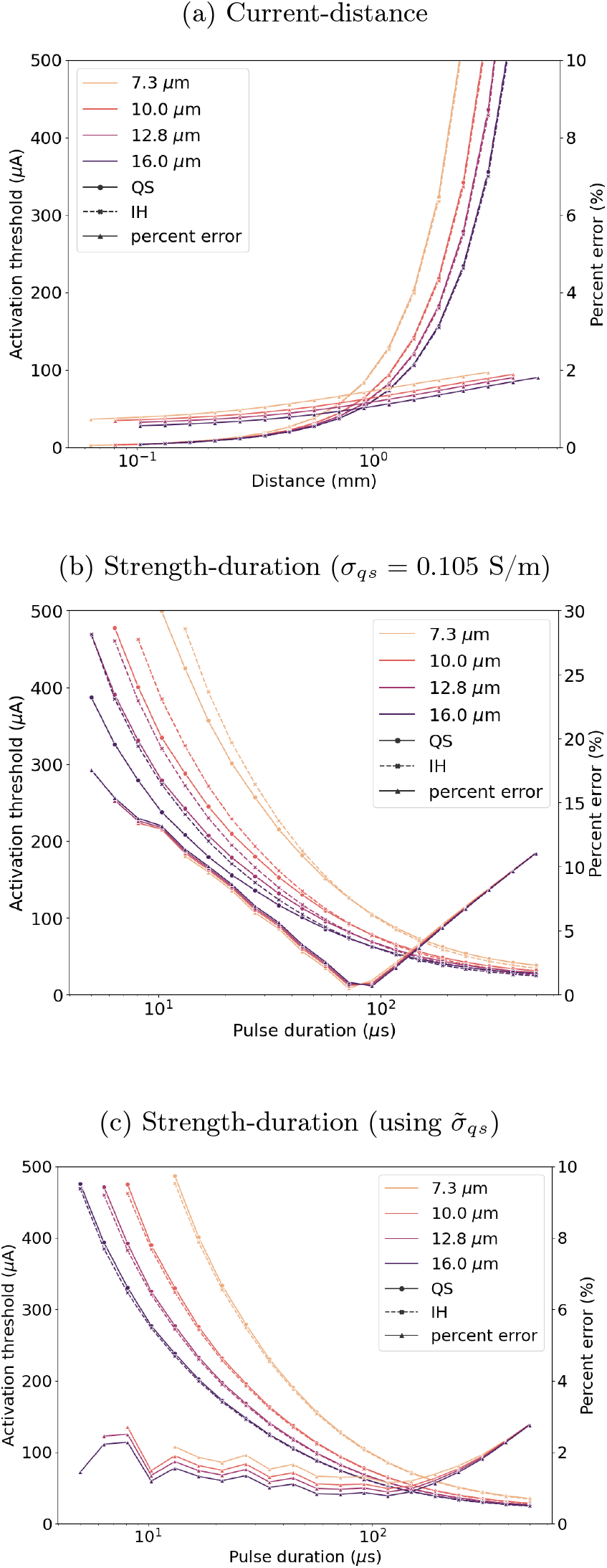
Activation thresholds calculated for QS and IH solutions for various fiber diameters. (a) Current-distance relationship, with a pulse duration of 100 *μ*s. (b)-(c) Strength-duration curve with a electrode-to-fiber distance of 1 mm. The conductivity for the QS approximation was *σ*_*qs*_ = 0.105 S/m in (a)-(b), and the value 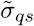 was calculated using (13) in (c).

Furthermore, in agreement with previous reports [29, 19], activation thresholds were reduced for larger fibers, and the errors between QS and IH solutions for both SD and IX curves appeared to be independent of fiber diameter.

### 4.3 Pulse train activation

We extended prior work [5] and quantified the errors in threshold when using the QS approximation for trains of pulses with different repetition rates and amplitudes.

We computed the fidelity for both field models and for different fiber diameters (figure 3a). We defined the cutoff frequency for a given amplitude as the lowest frequency at which fidelity dropped below 100%. No remarkable differences were observed across fiber diameters or models for the said critical frequency. In all cases, there was good agreement between QS and IH responses for stimulation frequencies *<* 700 Hz, for all fiber diameters, and we termed this region the *congruence area*. Yet, substantial differences were observed for frequencies above ≈700 Hz, *i*.*e*., where clear non-linear phenomena occured. Around 450 *μ*A and the 700 Hz – 1.5 kHz frequency range, a more dramatic fidelity loss is observed, which increases the fidelity error (from (16)) between models.

**Figure 3:**
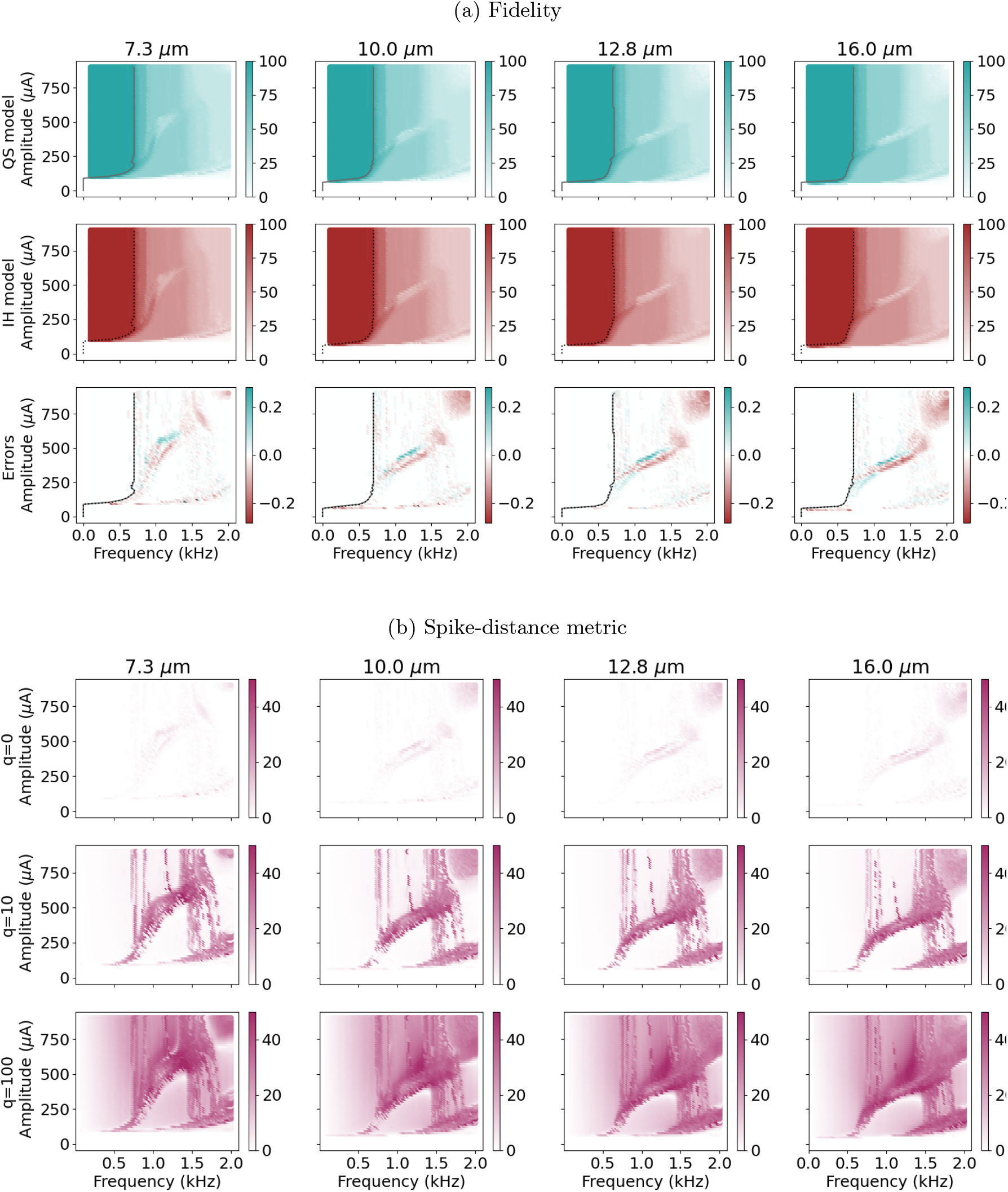
Action potentials generated by pulse trains across amplitudes (vertical axis), and repetition rates (horizontal axis), for different MRG fiber diameters (columns). (a) Fidelity index for the QS (top row, cyan) and IH (middle row, brown-red) models, and the fidelity relative error (bottom row). The grey lines indicate the cutoff frequency. (b) Spiketrain distance between the QS- and IH-generated action potentials, for parameter values *q* = 0, 10, 100 (top, middle, and bottom rows, respectively).

To quantify the shifts in time between QS and IH responses, we used the spike-train distance, calculated for different values of *q* (figure 3b). For *q* = 0 the spike-train distance was similar to the difference between fidelity maps for all frequencies, and, below 700 Hz the QS and IH models generated the same number of action potentials, . However, for higher values of *q* the spike-train distance increased monotonically in general, indicating consistent delays between trains, for all ranges of frequency. Still, outside the congruence area, we notice non-linear patterns, where trains generated by each model remain coherent or exhibit pronounced disparities in a complex manner.

To illustrate the differences between models, we generated raster plots for a small set of frequencies that evoked very different activation patterns (figure 4). We note the fidelity loss is clear from the number of delivered pulses versus the generated spikes with either model. Away from the congruence area the number of spikes was bounded because higher stimulation frequencies did not lead to a greater number of action potentials. As well, the timing differences in spikes generated by either model were evident but did not exhibit a linear relation with the frequency, in view of the fact that some action potentials were generated when using the IH equation instead of the QS approximation, or vice-versa.

**Figure 4:**
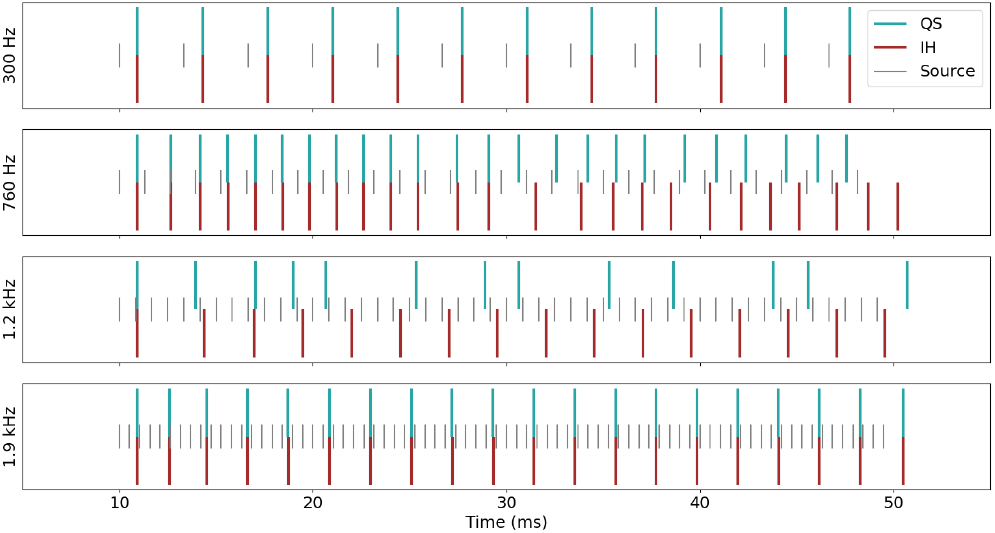
Action potentials generated by the QS, and IH models. Stimulation pulses from the source have amplitude *I* = 405 *μ*A.

**Figure 5:**
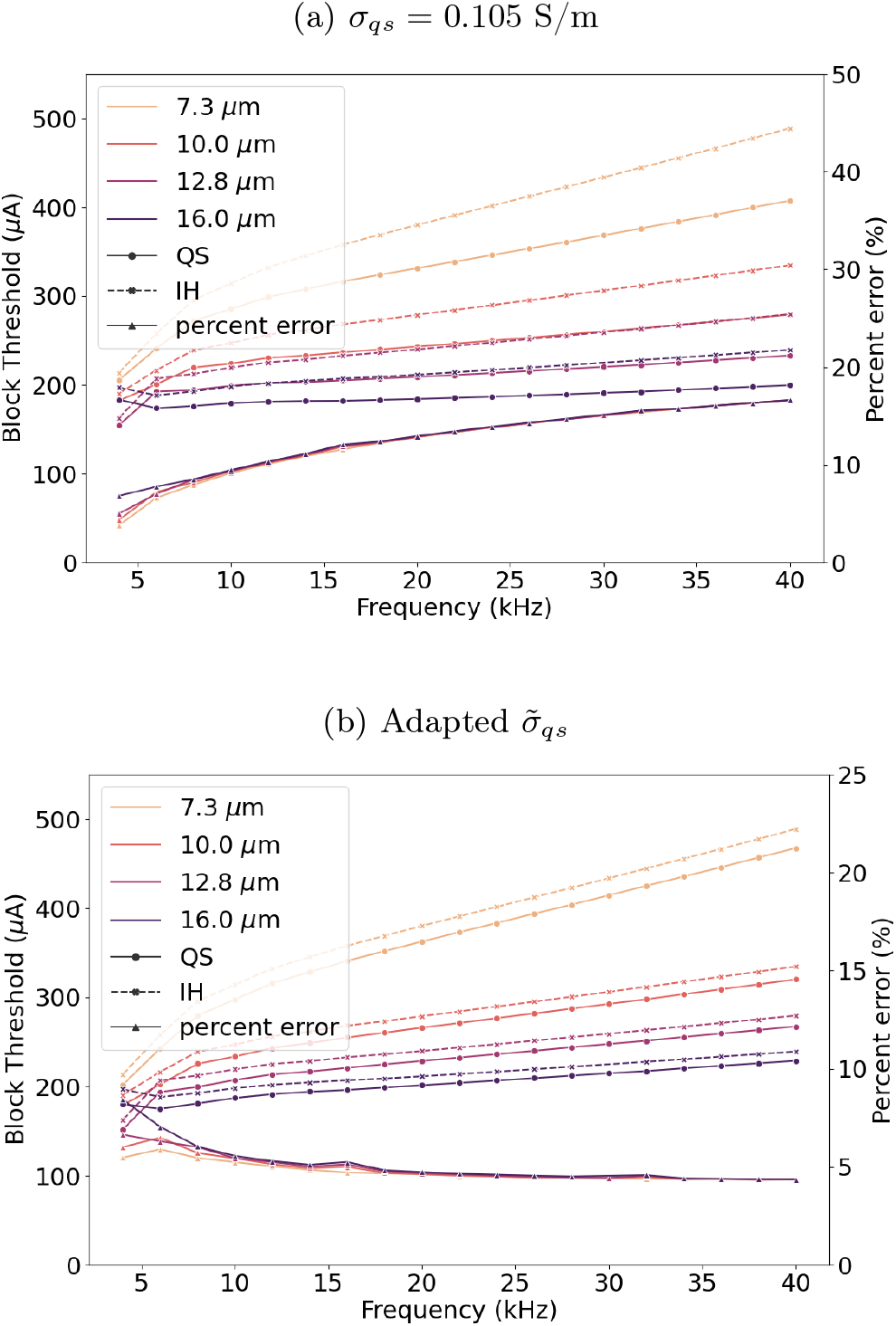
Conduction block with full-duty cycle pulses stimulation. Block thresholds were computed with the QS assumption (solid line) and the solution to the IH equation (dashed line). Relative errors are shown on the right axis. The effective conductivity for the QS electric potential is (a) fixed at *σ*_*qs*_ = 0.105 S/m, and (b) corrected to 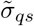, using (13)

### 4.4 Block thresholds

To quantify the differences in conduction block using the QS and the IH solutions, we applied the extracellular potential generated by either model. We replicated the setup from [3], where conduction block occurred near the center of the fiber if the block signal amplitude was high enough.

We set the pulse duration accordingly to the frequency of repetition, *i*.*e*., we use full-duty cycle pulse trains (recall the limit case for trains in (14)), and observed that block thresholds increased with frequency and were lower for larger fibers, in agreement with previous work [3]. For a fixed value of the conductivity, the slope of each curve was distinct, and errors increased from 3.8% to 16.6%. However, for the corrected quasi-static conductivity, the slope of the block threshold curve obtained with the QS model matched that of the IH model, Slight deviations were observed, and errors were limited to 8% overall, and for frequencies above 10 kHz reduced to less than 5%, whereas no remarkable differences were noted across fiber diameters.

For fixed-duration pulses, block thresholds decreased with increasing frequency. Choosing a naive value of the conductivity resulted in pronounced percent errors for frequencies *>* 10 kHz, of 17 and 20%, whereas for the corrected 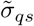, errors were reduced to 3-7%. However, substantial differences were observed between models across fiber diameters, even for corrected QS conductivities.

## 5 Discussion

The objective of this study was to determine under which conditions a more precise calculation of the electric potential is needed, when considering extracellular stimulation and block of model nerve fibers. Extending previous approaches that considered low signal frequencies [28], or limited the analysis to single pulses [5], the present analysis extended to trains of pulses with KHF repetition rates. A fixed value of the quasi-static conductivity at *σ*_*qs*_ = 0.105 S/m for grey brain matter provided valid calculations of the threshold for a pulse duration of 100 *μ*s, but errors up to 17% (figure 2b) appeared if the conductivity is not modified according with pulse duration. We introduced a power-based approach to revise the conductivity, showing that the QS approximation can be improved to obtain better accuracy for estimating activation thresholds with errors below 3% (see figure 2c) and block thresholds with limited errors to 5% for frequencies above 10 kHz (figure 5b). However, in this study, the scope of this correction is limited to energy-balanced potentials, such as biphasic pulse trains.

### 5.1 Capacitive effects and time delays

The most questionable simplification of the quasi-static assumption is presuming the media is purely resistive and neglecting the capacitive effects [28, 5, 22]. The capacitive effects can be neglected if the criterion *ωε/σ* ≈ 0 is satisfied. Still, even if all of the quasi-static assumptions are verified, errors up to 17% may occur, as observed in the SD curves (Figs. 2b and [5, figure 6a]). We observed reduced errors under 3% for appropriate values of the effective conductivity chosen through equation (13) because it allowed us to make a better approximation on the generated electric potential. Indeed, for short pulses, the capacitive effects damp the maximum amplitude reached by the IH model, so correcting the conductivity is in fact an adjustment to the QS amplitude, thereby transferring the same average power from the source to the fiber.

**Figure 6:**
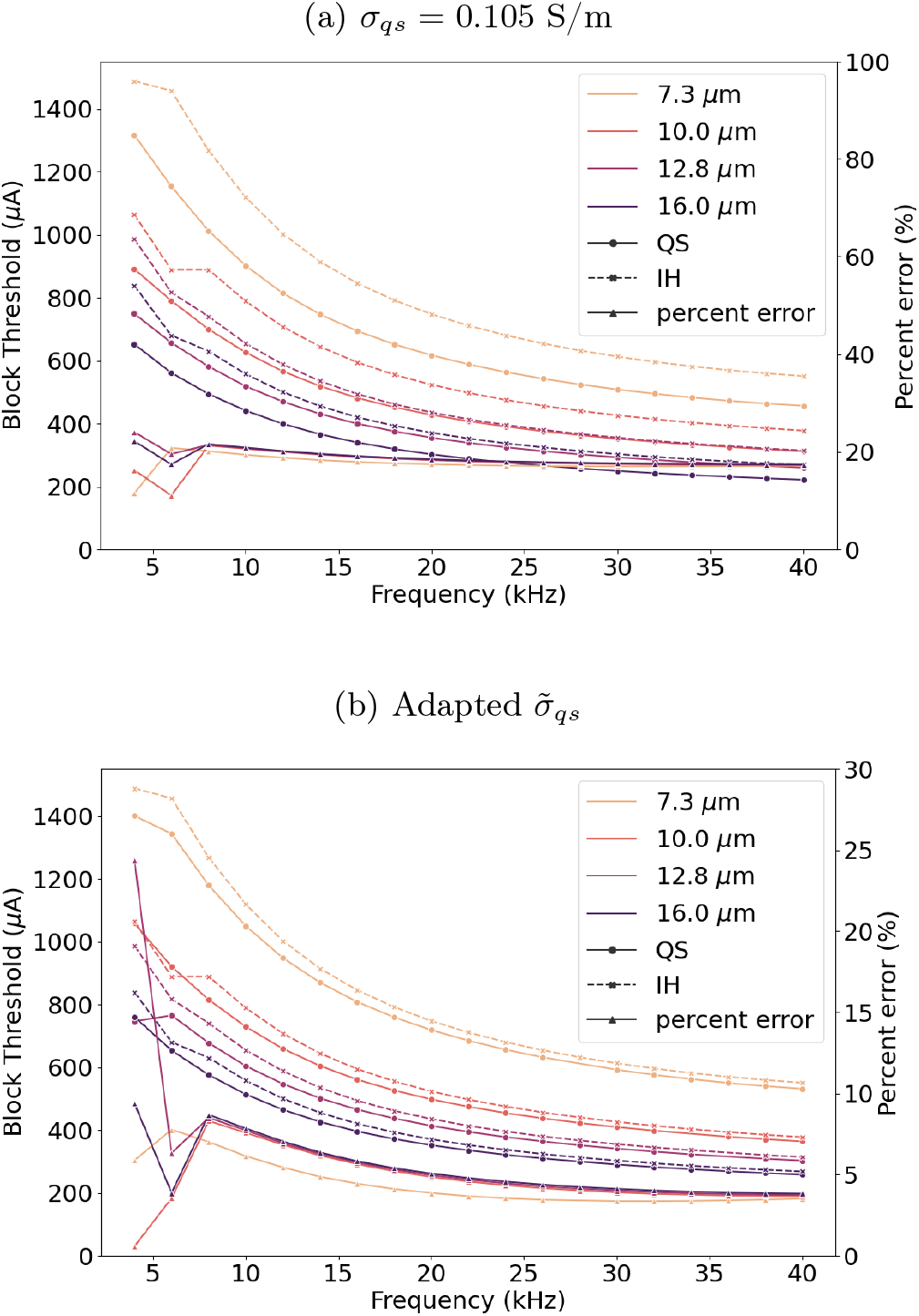
Conduction block threshold for trains of pulses of a fixed pulse duration of 10 *μ*s computed for stimulation under the QS assumption (solid line), and the solution of the IH equation (dashed line). The effective conductivity for the QS electric potential was (a) *σ*_*qs*_ = 0.105 S/m, and (b) corrected to 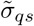, using (13). Percent errors are shown on the right axis.

Additionally, we found that as the frequency of stimulation increased, time shifts between the fiber responses became more substantial. The capacitive property of the medium prevents abrupt changes in the electric potential, and the dynamics of the resulting solution exhibit continuous charging and discharging, unlike the QS model, for which any discontinuity in the stimulation is directly reflected in the potential. Hence, reaching the threshold amplitude that triggers action potentials does not occur simultaneously with both models. This is quantified using high values of *q* in the spike-train metric, which expresses a measure of dissimilarity between trains. In figure 3b, there is a consistent shift in action potential timing, which can be explained by the transient effects neglected by the QS assumption.

### 5.2 Congruence region

We identified a congruence region, reflecting the range of validity of the QS assumption, for frequencies below 700 Hz. As shown in the fidelity maps (Figs. 3a), maximal fidelity was obtained only within this region for supra-threshold amplitudes. Near the threshold, the boundary of the congruence region was frequency-dependent, *e*.*g*., for 100 *μ*A fidelity dropped below 1 at about 100 Hz. This observation may be relevant for the analysis of, for example, novel sub-perception spinal cord modalities that may function in a regime of low fidelity despite using low (90 Hz) pulse repetition rates [20]. For frequencies above the cut-off frequency, the fidelity dropped dramatically, similar to experimental measurements in dorsal column axons [10]. Yet, for certain combinations of amplitudes and frequencies, the fidelity did not monotonically decay, but it exhibited a non-linear relationship with the repetition rate in small ranges.

Nevertheless, the spike-train distance showed that there were marked differences between the field models in the spiking activity within the congruence area (figure 3b). Even though fidelity was maximal, and the number of generated pulses was the same, as calculated for *q* = 0, for large values of *q*, the distance between spike trains increased, which is interpreted as a measure of growing dissimilarity (figure 3b). Therefore, the valid region of the QS assumption is contained within the congruence area but fails to cover the full extent of it. The spike-distance comparison was performed using a best-choice (according to [5]) effective conductivity, but still increased the penalty to shifts in time, *i*.*e*. costly *q*Δ*t*, emphasizes minor deviations in timings between models even for the classical range of frequencies. In addition, our findings suggest that synchronization of the firing activity to the stimulation signal is affected by the transient of the potentials, even for low pulse rates, *e*.*g* no more than 200 Hz, and this observation can be important in the design of electroceuticals devices, *e*.*g*., cochlear implants [30].

### 5.3 Conduction block

Conduction block using KHF signals can be achieved consistently with extracellular stimuli *>* 5 kHz [4]. Our results (figure 5a) show that the relative error between block thresholds calculated using the IH model vs the QS model with a fixed QS conductivity is not negligible, and increases steadily with the repetition frequency to more than 16%. We were able to reduce considerably inconsistency between the QS and the IH model from 16.6% to 5%, but the power-based argument appears to still be a partial correction due to residual errors for all frequencies. Nevertheless, prior work showed the threshold slopes using the QS assumption (with fixed conductivity) were less steep than *in vivo* measurements [27, 25], so choosing the corrected value using our method accounts for part of this discrepancy.

Prior work demonstrated that block thresholds decline for higher duty cycles for both symmetric and asymmetric rectangular pulse trains [27]. Accordingly, we observed that block thresholds decreased for higher pulse rates if the pulse duration was fixed at 10 *μ*s (figure 6). In this case, a higher frequency is equivalent to a lower duty cycle. Further, the error of the QS block threshold increased with increasing frequency for the full-duty cycle waveform, but, although larger, did not vary substantially for the fixed pulse duration waveform. This suggests that rather than the repetition rate, the error, in this case, is dominated by the frequency content of each pulse. Still, this argument holds for frequencies above 8 kHz in our results (recall figure 6b), or in general, remains valid provided the minimal frequency to generate consistent blocking is satisfied [25].

### 5.4 Effective conductivity

To compute the corrected QS conductivity (see equation (13)) we employed a novel method that determines the effective conductivity by matching the average power delivered from the source to the center of the fiber. We employed the definition of RMS voltage and current, that account for the direct values producing the same power in resistive loads, which is true under the QS assumptions, but not for dispersive media in general. Additionally, we note that this novel value is space-dependent, despite the initial assumption of isotropy, and should be computed separately for each fiber node. Consequently, the errors for activation and block thresholds of the QS model with corrected conductivity were not null, but were reduced to less than 1% and 3% respectively. This proposed framework can be extended to any energy-balanced waveform or living tissue (provided data are available), as illustrated in figure 7, yielding lower errors than those obtained with arbitrary values that are selected without considerations of the stimulus parameters.

**Figure 7:**
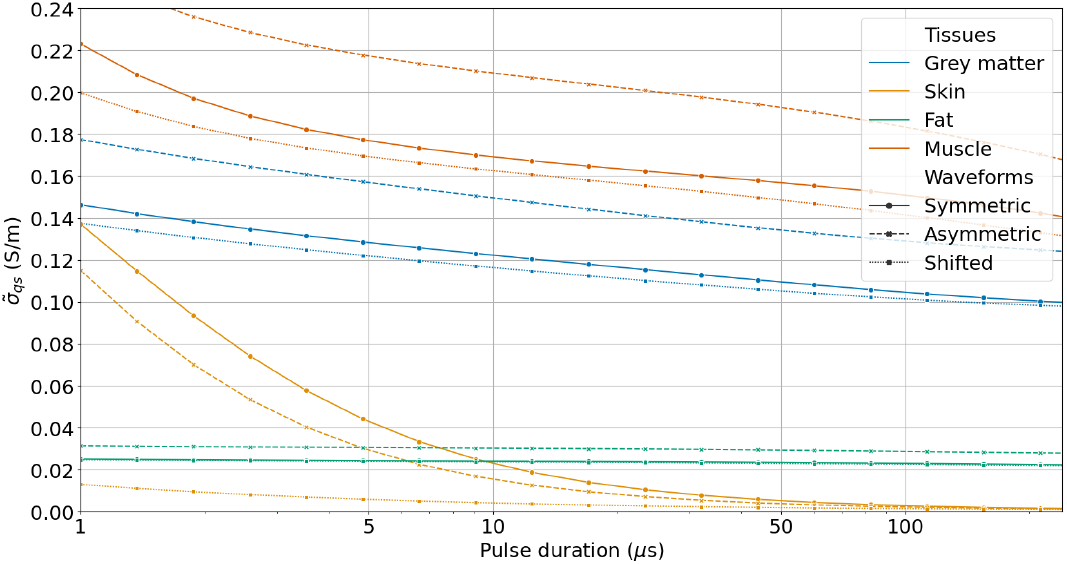
Corrected conductivity for the quasi-static solution, as a function of pulse duration (logarithmic scale). Effective conductivities for brain grey matter, (wet) skin, (not infiltrated) fat, and muscle are displayed, for sources with the waveforms defined in (17). Series truncated at 2 MHz.

Different signatures of effective conductivity for different tissues are directly linked to capacitive effects. For short pulses, the maximum amplitude of the potential is reduced, and thus the conductivity should be increased to reduce the amplitude of the QS solution, which is explicit in (10) because of the direct scaling. Notably, fat conductivity does not depend on the pulse duration, due to its nearly non-dispersive properties, but rather on the stimulation waveform itself.

### 5.5 Limitations

The computational methodology presented in this work has several limitations. First, we did not account for electrode shape and considered a simplified geometry. However, our motivation was to validate the mathematical assumptions rather than analyze a more complex model.

We assumed an infinite, homogeneous, and isotropic medium that does not reflect real tissues that exhibit finite sizes, anisotropy, and inhomogeneities. The linearity of the medium used to deduce the solution of the IH equation is a common assumption in neurostimulation [29, 26]. However, under certain conditions, the linearity of living tissues may break down, particularly if high-intensity electric fields are used [9]. Moreover, for computational purposes, we used truncated Fourier series to approximate the generated stimulus, giving rise to Gibbs phenomena in the time domain. To minimize this source of error, we included frequencies up to at least 50 times the frequency of repetition.

Lastly, the improved choice of conductivity in this work is limited to biphasic pulses generated by a point source. A rigorous and complete analysis of the proposed methodology in section 2.4 for more complex waveforms, dependence on the electrode geometry, and more general media assumptions is left as future work.

## 6 Conclusions

The quasi-static approximation is widely used in neurostimulation applications to simplify the calculation of the electric potential. We extended a previous analysis to quantify the error of this approximation for nerve activation and conduction block using trains of pulses with repetition rates of up to tens of kilohertz.

Our results indicate that the quasi-static assumption cannot be naively extended to repetitive stimulation with high-frequency content, and we determined the ranges for which the error may be negligible, *i*.*e*., a congruence area. Furthermore, we introduced a novel methodology to select a more appropriate value of conductivity for the quasi-static approximation, which can be extremely useful in neurostimulation modeling, especially for more recent applications that incorporate KHF signals.

## Supporting information

Effective Conductivity

## Data Availability

The code and data that support the findings of this study are available online at https://www.github.com/tcaussade/qs-khz-neurostimulation

## Author Contributions

LEM, MC, EC, and WMG conceived, supervised, and obtained funding and computational resources for the study. TC, EP, and LEM performed the experiments. TC, EP, LEM, and MC analyzed the data. TC, EP, and LEM wrote the manuscript. All authors revised, commented on, and approved the final version of the manuscript.

## Acknowledgements

This work was partially supported by ANID Millennium Science Initiative Program through Millennium Nucleus for Applied Control and Inverse Problems NCN19-161, a Research Exchange Award from the School of Engineering at Pontificia Universidad Católica de Chile, and NIH grants R37 NS040894 and R01 NS126376. Computational support as cluster access was provided by Duke University, Pontificia Universidad Católica de Chile, and Universidad de Santiago de Chile. We also thank Edgar Peña (Duke University) for his valuable comments on the block experiments, and Cristóbal Acosta (Universidad de Santiago de Chile) for technical assistance.

## Appendix Corrected conductivity

We introduced a novel method to set the effective conductivity used in the QS approximation, which primarily depends on the waveform shape and parameters, leading to the closed-form expression in (13). In this appendix, we demonstrate its versatility by determining numerical values for a variety of tissues and waveforms (figure 7). The tissue parameters were taken from [13] for brain grey matter, wet skin, not infiltrated fat, and muscle, then replaced in (5) to evaluate *ε*_*m*_ and *σ*_*m*_. In addition to the symmetric biphasic stimulation defined in (14), we also consider the *asymmetric biphasic* pulse train,

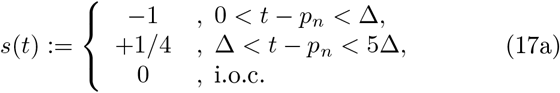

and the *shifted biphasic* pulse train,

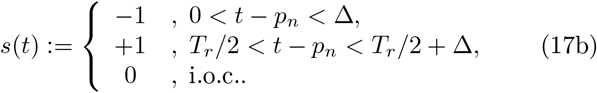

We considered the range of pulse duration for which the delivered power, energy, or charge to the fiber does not depend on the waveform [36] so that our proposed methodology is valid (figure 7). We observed that variations in the corrected conductivity were more significant for short pulses (*<* 50 *μ*s), where capacitive effects are more pronounced. For different waveforms, curves were qualitatively similar to a vertical shift, which resulted in remarkable differences for other tissues. We note that the effective conductivity of fat is constant, perhaps due to the small variation of conductivity of this tissue in this frequency range.

We recall the conductivity *σ*_*qs*_ = 0.105 S/m was introduced to minimize errors for 100 *μ*s pulses [5], which is consistent with our estimation and with the minimal

SD error (figure 2b). For pulses of 100 *μ*s, we calculated 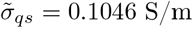 (with series truncated at 500 kHz), causing 4% difference with Bossetti et al. reference conductivity. In turn, such effective conductivity is obtained for a pulse duration of 94.468 *μ*s, producing a 5% error for the expected pulse duration.

