## Supplementary figures and images for "Towards a more accurate quasi-static approximation of the electric potential for neurostimulation with kilohertz-frequency sources^*^"

### Effective Conductivity

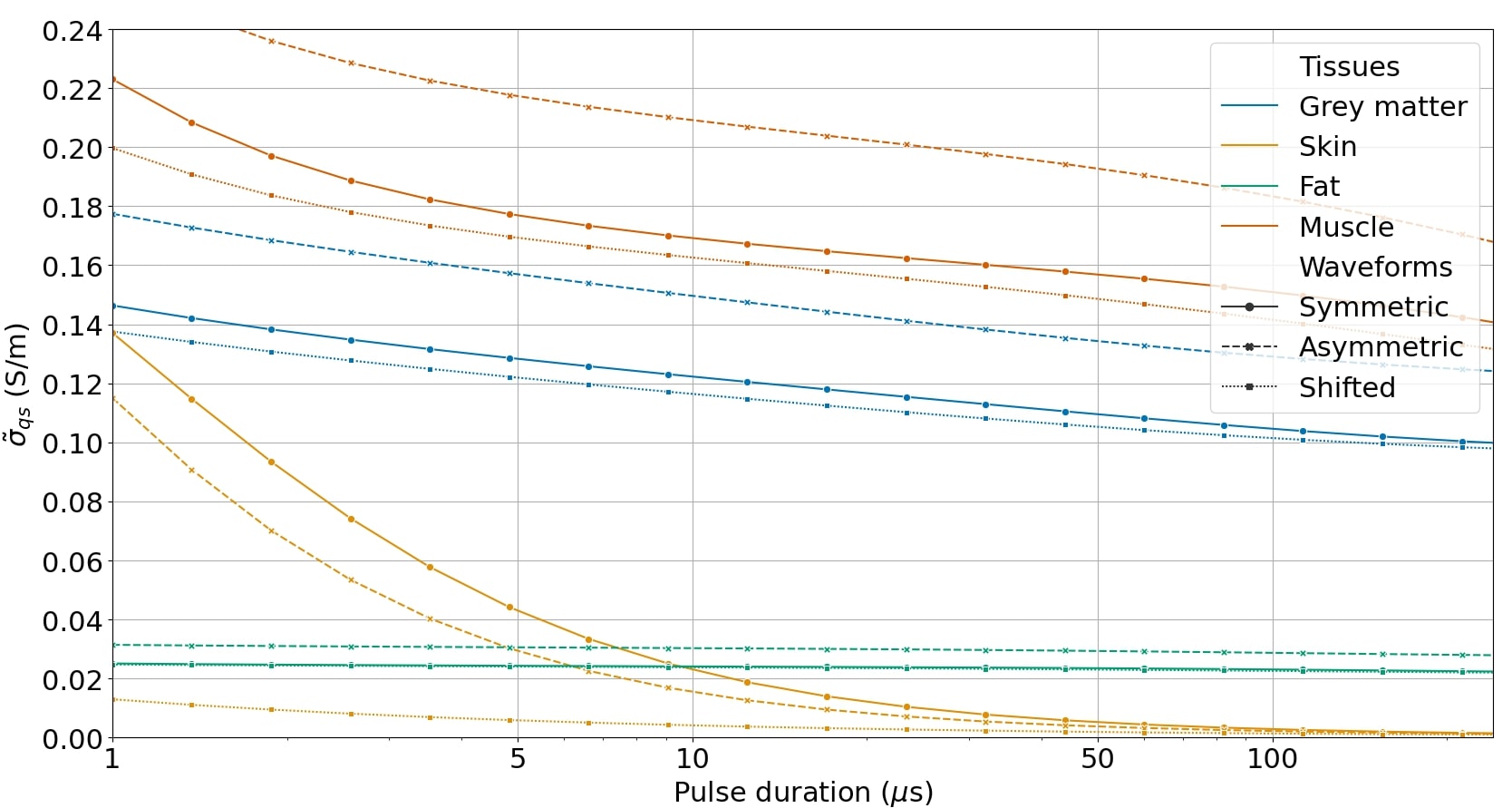
